# The Evolution of Human Ageing Under the Shadow of Demographic Transition

**DOI:** 10.64898/2025.12.03.691940

**Authors:** Handan Melike Dönertaş, Linda Partridge

## Abstract

The demographic transition has the potential to reshape the selective environment acting on the human genome. Here, we apply Hamilton’s force-of-selection framework to demographic schedules from 175 countries spanning 74 years (1950 to 2023). In post-transition populations, we observe an extension-dilution trade-off. The age at which selection intensity halves increased by 1.7 years since 1950, yet peak intensity declined by 29.4%. This decline was disproportionately severe at later ages. The ratio of selection intensity at age 20 to intensity at age 40 rose from 17.3 to 25.1, steepening the gradient favouring alleles with early benefits over late-life costs. Post-transition demography allows humans to function for decades beyond ancestral baselines, yet selection pressure to maintain late-life somatic integrity has never been weaker.

## Introduction

For most of human evolutionary history, our species lived under a consistent demographic regime marked by high mortality and correspondingly high fertility^1^. This environment shaped our genome, determining which alleles passed to the next generation and which deleterious mutations natural selection purged. However, the last century has witnessed a transformation without precedent in the history of human life. In just a few generations, a blink of an eye in evolutionary terms, we have overcome the demographic constraints that defined the selective environment of our ancestors^2,3^. This *demographic transition*, the shift from high birth and death rates to low birth and death rates as societies develop, has reshaped the human life course. Although the social and economic consequences of this transition are well documented, its evolutionary implications remain underexplored.

The evolutionary theory of ageing proposes that natural selection does not act uniformly across the lifespan. Instead, it acts strongly on early life but fades into a “shadow” as survival and reproductive potential decline^4^. This *selection shadow* creates a “biological blind spot” where late-acting deleterious mutations become less and less visible to natural selection and accumulate. Medawar proposed that selection weakens at advanced ages, leading to the accumulation of mutations with late-acting deleterious effects (mutation accumulation, MA)^5^. Williams extended this concept, recognising that alleles with early-life benefits can be favoured even if they impose late-life costs (antagonistic pleiotropy, AP)^6^. Crucially, these two mechanisms respond to different features of the selection curve: MA depends on absolute selection intensity at late ages, while AP depends on the gradient between early and late selection.

Hamilton mathematically formalised these ideas, deriving the intensity of selection across the lifespan^7^. He defined the force of selection at age 𝑥 as:

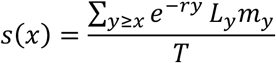

Here, 𝐿(𝑦) represents the person-years lived during age interval 𝑦, 𝑚(𝑦) denotes fertility at age 𝑦, 𝑟 is the population growth rate, and 𝑇 is the generation time. This metric integrates future reproductive contributions from age 𝑥 onward, weighted by survival and discounted by population growth. Hamilton’s framework predicts a specific shape for the selection curve: a constant plateau from birth until reproduction begins, a monotonic decline during reproductive years, and minimal selection beyond the age of last reproduction. Application of this framework to human populations revealed that selection acts in remarkably invariant ways across demographic diversity^8^; however, how 𝑠(𝑥) has changed *over time* as populations traverse the demographic transition remains unknown.

The demographic environment strictly determines the shape of the selection profile. In the last 200 years, global life expectancy has increased from approximately 25 to over 70 years^9,10^. Total fertility rates have declined from 6 to 7 births per woman to 1 to 2 births^11^. Compared with hunter-gatherer populations, infant mortality has fallen from 200 to 300 deaths per 1,000 births to fewer than 10^12^.

This transformation introduces a complex evolutionary trade-off. On one hand, falling mortality may extend the window during which selection operates. On the other hand, declining fertility and medical interventions may dilute the intensity of selection. The net result of these opposing forces, extension versus dilution, remains unknown.

We reconstructed the selection landscape of humans over the last 74 years. Using demographic data from 175 countries between 1950 and 2023, we calculated Hamilton’s 𝑠(𝑥) for the global population. We address three specific questions. First, how do survival, fertility and population growth combine to shape 𝑠(𝑥)? We use simulations to demonstrate how distinct demographic components mechanistically alter the selection shadow. Second, how much has 𝑠(𝑥) changed during the demographic transition? We characterise the magnitude and timing of these shifts to determine whether the selective window has expanded or contracted. Third, what are the evolutionary consequences for ageing? We examine how demographic change has reshaped the potential for both mutation accumulation and antagonistic pleiotropy, the two major mechanisms underlying the evolution of ageing.

## Results

### Demography Reshapes the Selection Curve

To isolate the independent effects of mortality, fertility and population growth on selection intensity 𝑠(𝑥), we modelled three synthetic demographic regimes (***Figure 1***). Scenario A represents a high-mortality, high-fertility regime (TFR = 7.6, 𝑒_0_ = 45.2) with early peak fertility. Scenario B represents moderate mortality with early-concentrated fertility (TFR = 2.5, 𝑒_0_ = 56), where reproduction peaks early and declines rapidly. Scenario C represents a low-mortality, low-fertility regime (TFR = 1, 𝑒_0_ = 82.7) with late peak fertility. We also developed an ***interactive simulation tool*** that allows continuous manipulation of these parameters (see Methods).

**Figure 1:**
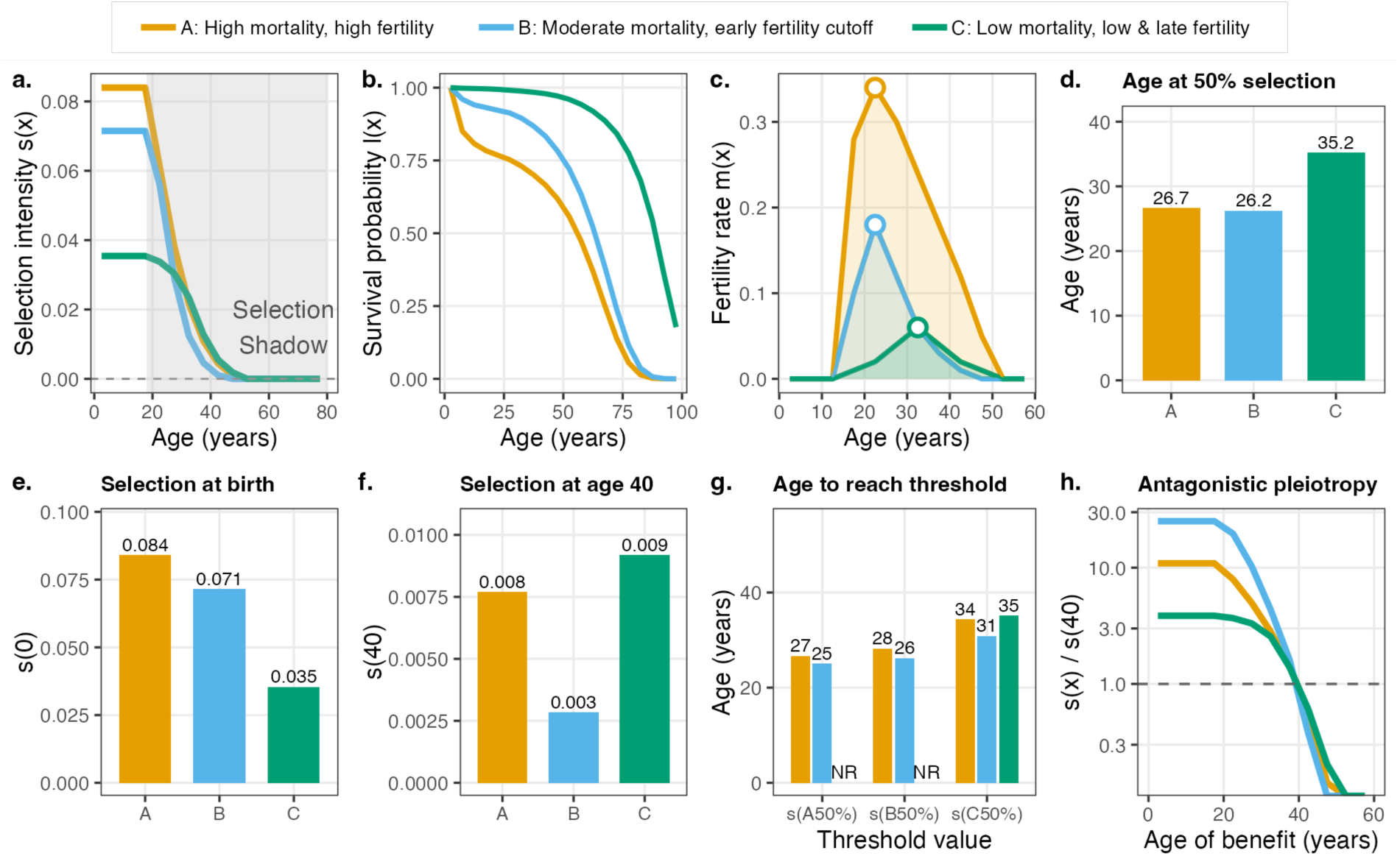
Selection curves across three demographic scenarios: A (orange; high mortality, high fertility), B (blue; moderate mortality, early-concentrated fertility), and C (green; low mortality, low/late fertility). **(a)** Selection intensity 𝑠(𝑥) versus age; grey region indicates the selection shadow where 𝑠(𝑥) declines. **(b)** Survival probability 𝑙(𝑥) versus age. **(c)** Fertility rate 𝑚(𝑥) versus age; circles (**O**) mark peak fertility, shaded areas represent cumulative fertility. **(d)** Age at 50% selection (age_50%_): the age when 𝑠(𝑥) drops to half its maximum. **(e)** Selection intensity at birth 𝑠(0) for each scenario. **(f)** Selection intensity at age 40, near the end of the reproductive window. **(g)** Age at which each scenario reaches specific selection thresholds (𝑠(A_50%_), 𝑠(B_50%_), 𝑠(C_50%_)); NR = never reached. **(h)** Antagonistic pleiotropy potential: ratio 𝑠(𝑥)/𝑠(40) versus age of early-life benefit; dashed line marks equal selection.

### Selection curves differ fundamentally across demographic regimes

Comparing different scenarios reveals distinct shapes of the selection curve (***Figure 1a***). Underlying shifts in demographic schedules drive these differences, where survival probability increases substantially (***Figure 1b***), while the fertility schedule varies in both timing and intensity (***Figure 1c***). Although all scenarios retain the theoretical pre-reproductive plateau, the intensity and trajectory of subsequent decline vary markedly. In Scenario A, selection collapses rapidly after reproduction begins, reaching 50% of its maximum intensity by age 26.7 (***Figure 1d***). Scenario B, with its early-concentrated fertility, shows the most rapid initial decline (age_’&%_ = 26.2), even faster than Scenario A, and continues to very low selection values due to its early fertility cutoff. In Scenario C, selection declines more gradually, extending the selection half-life to age 35.2.

### The extension-dilution trade-off

Despite the extended selective window in Scenario C, absolute selection force weakens. Selection intensity at birth 𝑠(0) declines by 58.33% from Scenario A (𝑠(0)=0.084) through Scenario B (𝑠(0)=0.071) to Scenario C (𝑠(0)=0.035) (***Figure 1e***). This delineates a fundamental evolutionary trade-off: shifting from high to low mortality and fertility regimes can extend the duration of selection but dilutes its intensity. Critically, selection at age 40 varies dramatically across scenarios: Scenario B shows the lowest value (𝑠(40)=0.003) due to its early fertility cutoff, while Scenarios A and C show higher values (𝑠(40)=0.008 and 0.009 respectively; ***Figure 1f***). The maximum selection intensity in Scenario C (𝑠(0)=0.035) never reaches the intensity that Scenarios A or B experience at their halfway points (𝑠(A_50%_)=0.042, 𝑠(B_50%_)=0.036; ***Figure 1g***).

### Predictions for ageing mechanisms

These simulations yield distinct predictions for the two pillars of evolutionary theories of ageing. For **mutation accumulation (MA)**, the critical parameter is absolute selection intensity at late ages. The decline in 𝑠(𝑥) from Scenario A to C predicts increased MA potential, as late-acting deleterious mutations face weaker purifying selection in low-mortality, low-fertility regimes. For **antagonistic pleiotropy (AP)**, the relevant parameter is the *gradient* between early and late selection. Here, the pattern is non-monotonic (***Figure 1h***). In Scenario A, selection pressure at age 20 is 9.1-fold higher than at age 40. Scenario B, despite lower absolute selection at age 20 (𝑠(20)=0.064 vs. 0.073), shows the *highest* gradient (21.3-fold) because its early fertility cutoff drives 𝑠(40) to very low levels (𝑠(40)=0.003). In Scenario C, the ratio decreases to 3.9-fold. This demonstrates that AP potential depends critically on the *shape* of the fertility schedule, not just overall fertility levels: early-concentrated reproduction can amplify the selection gradient even when absolute selection is moderate.

These simulations demonstrate that MA and AP respond to different features of the selection curve, namely absolute intensity versus relative gradient. Scenario B illustrates one route to increased AP potential, where early concentration of fertility drives s(40) toward very low levels, steepening the gradient. Similar effects can arise through other mechanisms, including the combination of fertility decline and timing shifts that leave minimal absolute reproductive potential at late ages. However, Scenario C shows that AP potential can also decline when late fertility is maintained alongside reduced overall selection. With multiple demographic parameters at play, the direction of change in AP potential is not straightforward. The actual evolutionary consequences in real populations will depend on how demographic transitions reshape both the level and timing of fertility, which we examine next.

### Selection Shadow Across Human Populations

#### Demographic transitions have reshaped the global selection curves

To quantify how evolutionary pressure varies across human demographic contexts, we analyzed a curated dataset of demographic schedules covering 175 countries and 16 aggregate regions from 1950 to the present (***Figure S 1***). We excluded country-years with populations below 100,000, extreme growth rates (|𝑟| > 10%) due to events such as mass migration and wars, and computationally implausible selection values (see Methods; **Tables S1–S3**). We integrated age-specific mortality, fertility, and population growth data to reconstruct selection intensity across countries (**Table S4**). This comprehensive panel allows us to apply Hamilton’s theoretical framework to empirical demographic data, revealing how the demographic transition has reshaped selection on late life. Between 1950 and 2019, the global mean population growth rate declined from 1.9% to 1.1% (***Figure S 2***), the global mean Total Fertility Rate (TFR) declined from 4.9 to 2.4 births per woman (***Figure S 3***), and life expectancy at birth (𝑒_&_) increased from 45.4 to 72.8 years, a gain of over 27 years in just seven decades (***Figure S 4***). These global averages, however, mask substantial regional heterogeneity. High-income countries showed decline in population growth while low-income regions maintained higher growth rates throughout (***Figure S 2***). High-income countries have sub-replacement fertility (𝑇𝐹𝑅_1950_=3, 𝑇𝐹𝑅_+2019_=1.6) and prolonged survival (𝑒_0_, 1950=61.8, 𝑒_0_, 2019=81.3). Low-income regions remain at an earlier stage of the demographic transition, with higher fertility and lower survival (𝑇𝐹𝑅_1950_=6.5, 𝑇𝐹𝑅_+2019_=4.8, 𝑒_0_, 1950=30.3, 𝑒_0_, 2019=63) (***Figure S 3, Figure S 5***). This variation produces divergent selection landscapes across regions (***Figure 2, Figure S 6***). In pre-transition contexts (e.g., Niger; ***Figure 2a***), selection intensity shows a high initial plateau followed by a steep decline, reflecting high mortality and early fertility. In contrast, post-transition contexts (e.g., Japan; ***Figure 2a***) exhibit lower baseline intensity (𝑠(0)) but a more prolonged decline, driven by the rectangularization of the survival curve (***Figure 2b***) and the delay of reproduction (***Figure 2c***). In 1950, Japan’s selection curve resembled pre-transition profiles (𝑠(0)_Japan,1950_=0.064, 𝑠(0)_Niger,1950_=0.061). By 2019, baseline selection intensity had declined by 31.3% in Japan while increasing by 27.9% in Niger (𝑠(0)_Japan,1950_=0.044, 𝑠(0)_Niger,2019_=0.078).

**Figure 2:**
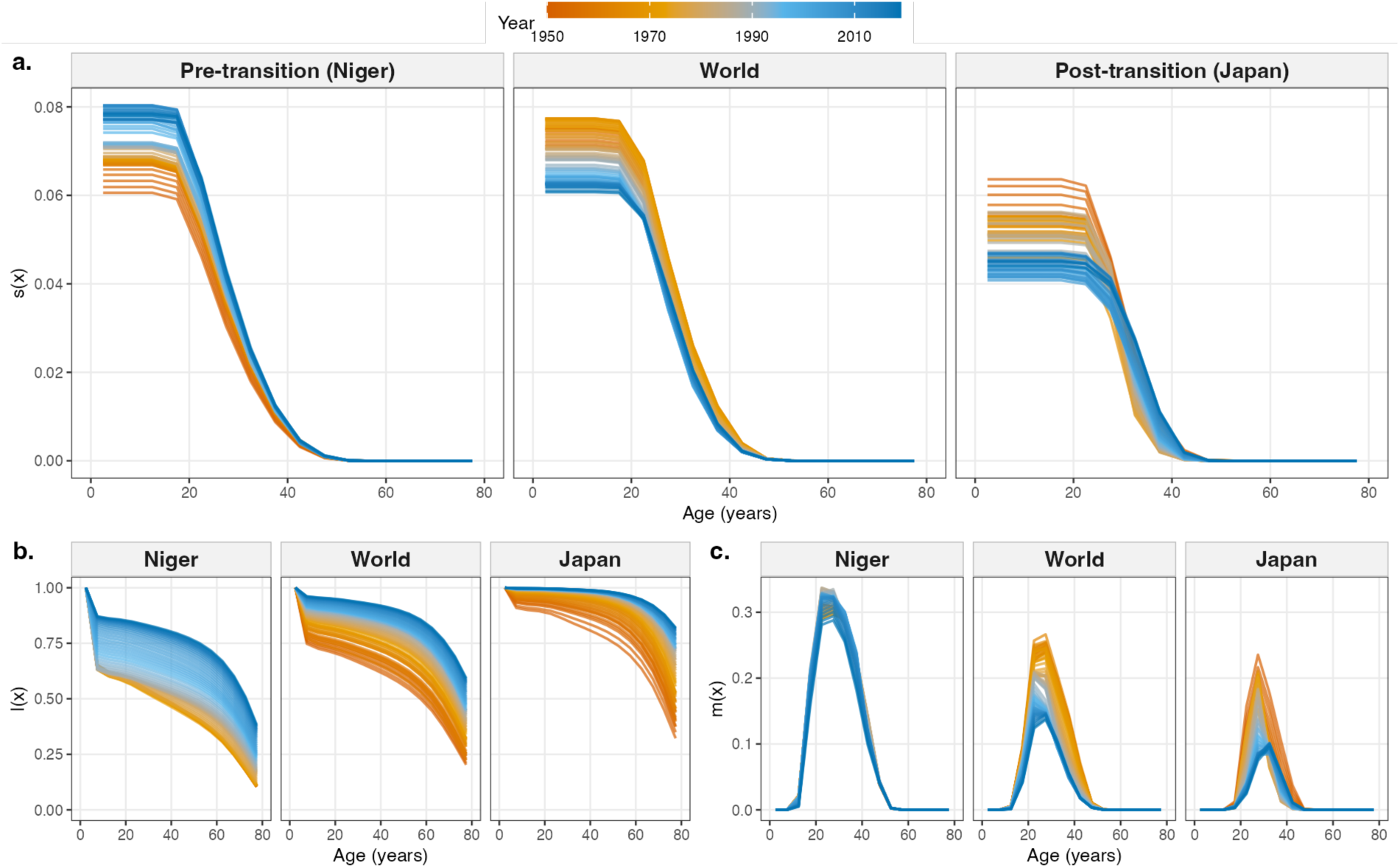
Selection curve varies by demographic context. **(a)** Selection intensity 𝑠(𝑥) for three representative cases: Niger (pre-transition), World, and Japan (post-transition). Curves are colored by year from 1950 (orange) to 2019 (blue). **(b)** Survival probability 𝑙(𝑥) showing rectangularization of mortality in post-transition countries. **(c)** Fertility rate 𝑚(𝑥) demonstrating both the decline in peak rates and the shift toward later fertility.

### Temporal Shifts in Selection Landscape

The global surface of selection intensity has shifted systematically over the past 74 years. This “extension-dilution” dynamic is pervasive. A comprehensive analysis of 175 countries confirms that while the window of selection has widened globally, its maximum force has weakened in most contexts (***Figure 2, Figure S 6***). High-fertility countries generally retain higher 𝑠(0) values, whereas post-transition low-fertility, low-mortality countries cluster at lower intensities with more gradual declines.

### Selection trajectories diverge by demographic stage

The global average masks a profound polarization in evolutionary trajectories driven by the pace of demographic transition (***Figure 3*, Table S5**). Countries retaining high fertility (𝑇𝐹𝑅 > 4, n=36) exhibit stable selection landscapes. Their selection profiles have remained largely unchanged over 74 years (***Figure 3a***), showing no significant directional trend in selection at birth (𝑠(0)) (median 𝜌 = 0.11, *p*=0.783) or the selection window age_’&%_ (median 𝜌 = 0.25, *p*=0.342, ***Figure 3c-d***). Post-transition countries (𝑇𝐹𝑅 ≤ 2, n=77) have undergone a collapse in selective potential (***Figure 3a***). Selection at birth (𝑠(0)) declined (median 𝜌 =-0.71, *p*=2.9×10^-11^), falling from a mean of 0.072 in the 1950s to 0.051 in the 2010s. Simultaneously, the relative selective window has expanded, with the half-life of selection (age_50%_) extending by nearly 1.7 years (median 𝜌 = 0.48, *p*=2.2×10^-6^, ***Figure 3d***). Countries currently in transition (2 < 𝑇𝐹𝑅 ≤ 4, n=62) display intermediate patterns, with declining selection at birth (𝑠(0)) (median 𝜌 =-0.48, *p*=1.1×10^-6^, ***Figure 3c***) but no significant change in the selection window age_50%_ (median 𝜌 =-0.23, *p*=0.08, ***Figure 3d***).

**Figure 3:**
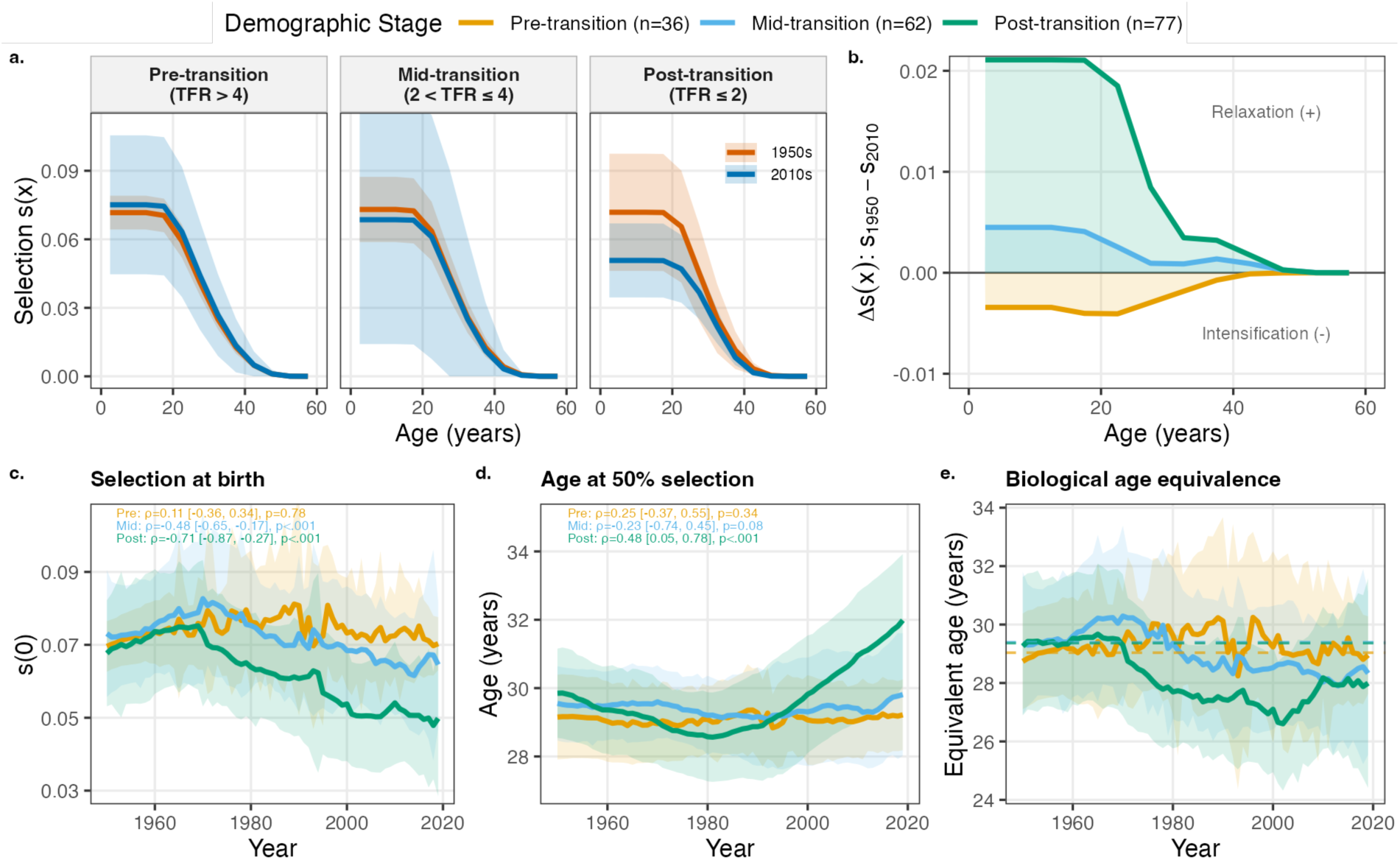
Divergent evolutionary trajectories by demographic stage. **(a)** Mean selection intensity 𝑠(𝑥) for Pre-(orange), Mid-(blue), and Post-transition (green) countries, comparing the 1950s (orange line/ribbon) to the 2010s (blue line/ribbon). Ribbons indicate ±1 SD. **(b)** Age-specific change in selection intensity (𝛥𝑠(𝑥)) between 1950 and 2010. Positive values indicate relaxation (weakening selection); negative values indicate intensification. **(c, d)** Temporal trends in **(c)** selection intensity at birth 𝑠(0) and **(d)** the age at 50% selection. Inset annotations report within-country temporal trends. For each country, Spearman’s 𝜌 was calculated between the metric and calendar year; the median 𝜌 across countries within each demographic stage is shown with the interquartile range [IQR], and p-values are from Wilcoxon signed-rank tests assessing whether the distribution of country-level correlations differs from zero. **(e)** Biological age equivalence, defined as the age at which a country experiences the selection intensity (𝑠(𝑥)) that was characteristic of the stage-specific “selection half-life” in the 1950s (dashed lines). The decline in the post-transition line indicates that comparable levels of selection are now restricted to younger ages.

### Selection relaxed in post-and mid-transition countries while intensifying in pre-transition countries

The differential 𝛥𝑠(𝑥) between the 1950s and 2010s reveals opposing dynamics across demographic stages (***Figure 3b***). Post-and mid-transition countries show relaxation (positive 𝛥𝑠) across all ages, with the greatest absolute loss of selective pressure occurring in early adulthood (ages 15 to 30) in post-transition countries. Pre-transition countries show intensification (negative 𝛥𝑠) throughout the age range, driven by improvements in childhood survival that increase the probability of reaching reproductive age and thus raise selection intensity at all ages. While the window of selection has widened in modern contexts due to longer lifespans, the absolute evolutionary value of any given year of life has been diluted. We quantified this dilution by calculating biological age equivalence, defined as the age at which a modern individual faces the same selection pressure that defined the selection half-life in the 1950s (***Figure 3e***). In post-transition countries, this equivalent age has dropped from 29.7 years to 28 years. Notably, while the selection half-life shows a monotonic trend, the tail of the selection shadow (ages at 10%, 5%, and 1% thresholds) exhibits a distinct U-shaped trajectory in post-transition countries, initially contracting before expanding in recent decades (***Figure S 7***).

### Demographic determinants of selection strengthen over time

The relationship between demographic parameters and selection has tightened over time (***Figure S 8, Figure S 9***). Bivariate correlations strengthened: the TFR-𝑠(0) correlation increased from 𝜌 = 0.25 to 𝜌 = 0.63, while the e₀-𝑠(0) correlation shifted from 𝜌 =-0.09 to 𝜌 =-0.67. While fertility shows the strongest association in multivariate models (***Figure S 9*d**), the high correlation among predictors makes attributing independent effects difficult.

### Implications for Evolutionary Theories of Ageing

The **mutation accumulation (MA)** hypothesis predicts that mutations with deleterious effects confined to late ages will escape purifying selection when selection intensity is low. ***Figure 4a*** shows striking differences across demographic stages. Pre-transition populations maintain consistently high selection intensity at young ages (orange region) with relatively stable contours over time. In contrast, post-transition populations show a pronounced temporal decline; the contour lines shift systematically toward younger ages, indicating that the same level of selective pressure now operates at younger ages. This creates an expanded “blind spot” where late-acting mutations face reduced purifying pressure, with the effect most dramatic in post-transition populations.

**Figure 4:**
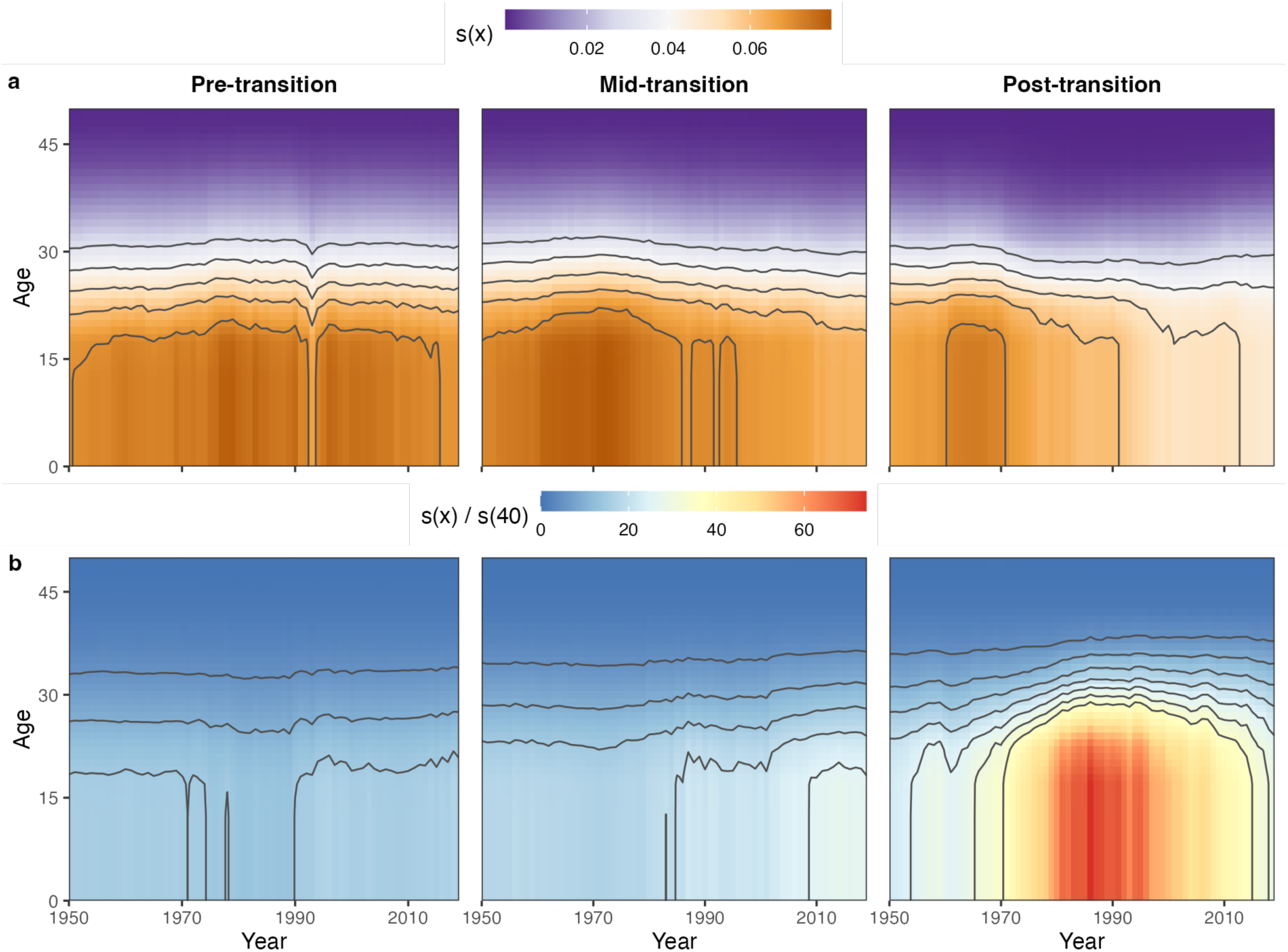
Implications for ageing mechanisms across demographic stages. **(a)** Lexis surfaces of absolute selection intensity 𝑠(𝑥) for Pre-, Mid-, and Post-transition populations. Orange indicates high selection intensity; purple indicates low intensity. Contour lines mark constant selection values. The temporal gradient from high (left) to low (right) selection is most pronounced in post-transition populations. **(b)** Selection gradient surfaces showing the ratio 𝑠(𝑥)/𝑠(40) across ages and time. Higher values (red/orange) indicate stronger selection at age 𝑥 relative to age 40. This ratio quantifies the potential for antagonistic pleiotropy, capturing the selective advantage of early-benefit alleles relative to their late-life costs. The dramatic intensification of the gradient in post-transition populations (right panel) reveals a steepening gap between early and late selection.

The **antagonistic pleiotropy (AP)** hypothesis depends not on absolute selection but on the *gradient* between early and late ages. Alleles conferring early benefits can spread even if they impose late costs, provided the early-age selection advantage exceeds the late-age cost. We quantified this gradient as the ratio 𝑠(𝑥)/𝑠(40) (***Figure 4b***). The contrast across demographic stages is remarkable. Pre-and mid-transition populations show relatively flat gradient surfaces (blue tones), indicating a lower gap between early and late selection. Post-transition populations, however, display an intensification of the gradient (orange/red region emerging after 1970), particularly at ages 15 to 30. At age 20, this ratio increased from 17.3 in 1950 to 25.1 in 2019, a 45% increase. This steepening gradient suggests that AP alleles face *stronger* relative selection in modern post-transition populations, as early-life benefits are now proportionally more advantageous compared with late-life costs. Paradoxically, while absolute selection has weakened everywhere, the gap between early and late selection has widened, potentially intensifying selection for AP variants in post-transition populations.

## Discussion

### The extension-dilution trade-off

A central theme emerging from our analysis is the extension-dilution trade-off, where falling mortality extends the selective window (age_’&%_ increases) while fertility decline dilutes intensity (𝑠_&_ decreases). Overall, dilution dominates, and selection weakens despite the longer selective window. The maximum selection intensity in post-transition populations falls below the half-life intensity of pre-transition populations, meaning a 28-year-old today faces selection pressure comparable to a 30-year-old in the 1950s. This complements prior work showing that demographic transitions alter variance in fitness and selection gradients on specific traits^13–16^, by revealing that the demographic transition restructures not just trait-specific selection but the entire age-specific selection landscape. The result is a fundamental evolutionary mismatch^17^. We have extended the need for somatic maintenance but weakened the evolutionary incentive to maintain it.

### Divergent trajectories

The demographic transition has created a polarisation in evolutionary trajectories. Pre-transition populations exhibit stability, with their selection landscapes remaining largely unchanged and childhood survival improvements actually *intensifying* early-age selection.

High extrinsic mortality creates selection bias toward early life^18^, and preindustrial populations showed high selection intensity where survival and fertility accounted for most fitness variance^19^. Post-transition populations show collapse, where 𝑠_&_ has declined precipitously while the selective window has extended, paralleling documented microevolutionary responses during demographic transitions^20^. Mid-transition populations display intermediate dynamics. This divergence means that human populations across different countries now operate under fundamentally different selective regimes.

### Implications for evolution of ageing

Both major mechanisms of evolution of ageing, mutation accumulation (MA) and antagonistic pleiotropy (AP)^5,6^, show amplified potential under post-transition demography. MA potential increases because absolute selection at late ages has weakened. Late-acting deleterious mutations would face reduced purifying pressure, consistent with evidence that the phenotypic variance attributable to genetic factors increases at older ages^21^. AP potential also increases, but through a different mechanism. Our simulations demonstrated that change in AP potential may increase (e.g., Scenario B with early-concentrated fertility) or decrease (e.g., Scenario C with late fertility) under lower fertility regimes, and the result depends on the curve shaped by interaction between fertility and mortality and their schedules. In empirical data, post-transition populations show the former pattern: the disproportionate collapse of selection at late ages steepens the early-to-late gradient. Our data show that the ratio 𝑠(𝑥)/𝑠(40) has increased substantially at early reproductive ages in post-transition populations. This steeper gradient means that alleles conferring early benefits would face proportionally stronger selection relative to their late-life costs. Existing genomic evidence supports both MA and AP in human ageing^22^, while age-related diseases cluster by onset profiles with distinct evolutionary signatures^23^. Whether the recent demographic shifts documented here will eventually contribute to such patterns remains a question for future investigation.

### Limitations

Several limitations should be considered when interpreting our findings. i) *Methodological assumptions.* Our calculations assume stable population dynamics. We excluded observations where extreme growth rates from wars or mass migrations violate this assumption, but instantaneous growth rates remain imperfect proxies for long-term fitness. Additionally, our stage classification based on 2019 TFR does not capture within-period transitions, meaning countries that shifted between stages during the study period are classified by their endpoint only. Finally, we restricted analysis to 1950 to 2019 to avoid COVID-19 disruptions and excluded small populations (fewer than 100,000) and biologically implausible extreme values to ensure data quality. ii) *Upper-bound estimates*: Our estimates represent an *upper bound* on true selection intensity. Medical interventions decouple phenotype from fitness in ways our demographic metrics cannot capture. Examples include obstetric interventions that allow survival of mother-infant pairs who would not have survived historically^24^ and treatments for conditions such as type 1 diabetes that would otherwise be lethal before reproduction^25^. Realised relaxation of selection is almost certainly greater than our calculations suggest. iii) *Scope of inference*: We measure demographic *potential* for selection, not genomic outcomes, and whether this relaxation manifests in changing allele frequencies in evolutionarily relevant timescales requires direct genomic investigation. Our female-centric fertility analysis may obscure sex-specific patterns in reproductive variance. Finally, Hamilton’s 𝑠(𝑥) captures sensitivity of fitness to survival changes but does not account for kin selection or grandparental effects that may maintain selection at post-reproductive ages in limited circumstances^26,27^, although empirical evidence suggests these indirect fitness contributions diminish substantially at advanced ages^28^.

## Conclusion

We have quantified the demographic determinants of the selection shadow across 70 years of global demographic change. The demographic transition emerges not merely as a socioeconomic transformation but as an evolutionary reshaping of the selective landscape acting on the human genome. As humanity converges toward low fertility and long life, this selection shadow may continue to expand. Such relaxation of selection could, over evolutionary timescales, lead to gradual genetic changes affecting baseline fitness^29^, though the timescales and magnitude remain debated. If late-acting deleterious mutations accumulate under relaxed selection, they could eventually contribute to the burden of chronic diseases that dominate modern morbidity, although detecting such effects would require genomic data spanning many generations, a timescale that likely exceeds practical observation. Future research must bridge the gap between these demographic potentials and genomic reality, determining whether relaxation is already visible in the changing frequencies of age-dependent alleles in human populations.

## Methods Calculations

### Life Table Construction

Survival schedules were calculated as 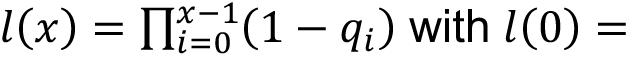 1. Person-years lived used linear approximation: 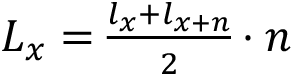 for closed intervals (𝑛 = 5), and 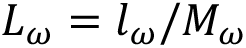 for the open interval, with a minimum threshold on 𝑀_ω_ to ensure numerical stability.

### Evolutionary Metrics

Net reproductive rate was calculated as 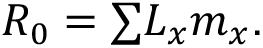 Generation time, discounted by the population growth rate (𝑟), was computed as 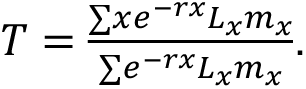

### Hamilton’s Selection Shadow

Following Hamilton^7^, the force of selection was calculated as 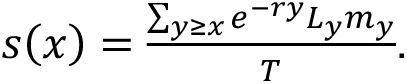 We treated 𝑠(𝑥) as continuous via linear interpolation to determine threshold ages (e.g., where 𝑠_x_ < 0.01) through inverse interpolation.

### Selection Metrics

Baseline intensity 𝑠_0_ measures selection at birth. The selection half-life (age_50%_) is defined as the age when 𝑠(𝑥) = 0.5 ⋅ 𝑠_0_. Selective tail thresholds (age_50%_, age_1%_) mark ages where selection falls below 5% and 1% of baseline. The area under the curve (AUC) represents total accumulated selection across the lifespan.

### Validation

All observations passed two automated checks: (i) identity validation (𝑠(0) × 𝑇 = 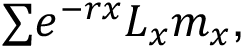, and (ii) shape validation confirming pre-reproductive plateau (CV < 5% for ages < 15), monotonic decline (ages 15–50), and post-reproductive convergence to zero.

### Demographic Simulation

We constructed three synthetic regimes to isolate theoretical effects of mortality, fertility, and growth on selection intensity.

### Schedule Construction

Synthetic life tables used 5-year intervals from birth to 100+ (21 age groups). Age-specific 𝑞_x_ and 𝑚_x_ were manually specified; all scenarios share a reproductive window (ages 15–49) but differ in mortality, fertility magnitude, and timing.

### Scenario Descriptions

Scenario A (high mortality, high fertility): 𝑞_0_ = 0.15, 𝑇𝐹𝑅 = 7.6 (peak 20–24), 𝑟 = 3.5%, 𝑒_0_ = 45.2 years. Scenario B (moderate mortality, early fertility peak): 𝑞_0_ = 0.04, 𝑇𝐹𝑅 = 2.5 (peak 20–24), 𝑟 = 1%, 𝑒_0_ = 56 years. Scenario C (low mortality, late fertility): 𝑞_0_ = 0.002, 𝑇𝐹𝑅 = 1 (peak 30–34), 𝑟 =-0.5%, 𝑒_0_ = 82.7 years.

### Selection Analysis

For each scenario, we calculated 𝑠(𝑥) and derived age_’&%_ via interpolation. We quantified extension-dilution by comparing 𝑠(𝑥) at fixed ages and cross-scenario threshold ages.

### Interactive Application

An open-access web application (R Shiny compiled to WebAssembly via shinylive) enables exploration beyond discrete scenarios. Mortality follows a Gompertz function 𝜇(𝑥) = 𝛼𝑒^βx^; fertility follows a Gaussian 𝑚(𝑥) = 𝑚_peak_ exp (-(x-𝜇_age_)^2^/2𝜎^2^, with 𝜎 bounding reproduction within ±3𝜎.

## Data Acquisition

### Sources

Age-specific demographic schedules were obtained from Our World in Data (OWID), including population by age and sex, birth counts, death counts, and growth rates (𝑟). The temporal scope spans 1950 to present, however, throughout we show mainly comparisons between 1950s (1950-1959) and 2010s (2010-2019) to ensure stable demographic patterns by excluding the COVID-19 pandemic.

### Variable Construction

Data were harmonised to 5-year intervals (0 − 4,…,100 +). Fertility rates were calculated as 𝑚_x_ = births / female population, with missing values set to zero for non-reproductive ages. Mortality rates 𝑀_x_ = deaths / population were converted to probabilities using 𝑞_x_ = 1 – 𝑒^-5mx^, with 𝑞_100+_ set to 1.0.

### Exclusion Criteria

(1) Small populations (< 100,000): removed 62 entities (3,433 country-years; **Table S1**). (2) Extreme growth (|𝑟| > 10%): because our selection shadow calculation relies on the stable population assumption, where age structure is determined by stable vital rates, we excluded 37 country-years across 17 countries reflecting structural shocks where instantaneous growth rates do not reflect biological fitness (**Table S2**). (3) Values that have unrealistic selection values (𝑠(0) ≤ 0 or ≥ 0.20) were removed 53 country-years (**Table S3**).

## Stratification and Temporal Analysis

### Temporal Aggregation

Country-year metrics were aggregated to decadal means. The extension-dilution trade-off was quantified as relative change in 𝑠_0_ and absolute shift in age_50%_ between the 1950s and 2010s.

### Demographic Stages

Countries were classified based on 2019 TFR into three stages: pre-transition (𝑇𝐹𝑅 > 4), mid-transition (2 < 𝑇𝐹𝑅 ≤ 4), and post-transition (𝑇𝐹𝑅 ≤ 2).

### Temporal Dynamics

Relaxation intensity was calculated as 𝛥𝑠(𝑥) = 𝑠(𝑥)_1950s_ − 𝑠(𝑥)_2010s_, where positive values indicate relaxation (weakening of selection). Within-country temporal trends were assessed using a two-step procedure to avoid pseudoreplication from repeated country-year observations. First, for each country, we calculated the Spearman correlation (𝜌) between each selection metric (e.g., 𝑠(0), age_50%_) and calendar year using all available annual data. Second, we summarized the distribution of these country-specific correlations within each demographic stage using the median and interquartile range (IQR). Statistical significance was assessed using a Wilcoxon signed-rank test to determine whether the distribution of country-level correlations differed from zero. This approach appropriately accounts for the hierarchical structure of the data where countries are the independent units. Biological age equivalence measures the age at which a modern (2010s) population experiences the same selection intensity that defined the half-life (age_’&%_) of the 1950s; a decrease indicates that comparable evolutionary pressure is now restricted to younger ages. Correlation matrices (Spearman) were computed between selection metrics and vital rates for the 1950s versus 2010s.

### Lexis Surfaces

Selection intensity 𝑠(𝑥) and the AP ratio (𝑠(𝑥)/𝑠(40)) were visualised across age × year. Stage-stratified analyses used median aggregation for outlier robustness, and surfaces were interpolated to 1-year resolution in both dimensions.

## Statistical Modelling

We performed standardised multivariate regression on decadal means (1950s, 2010s). Selection metrics (𝑠(0), age_50%_, age_5%_) were modelled as functions of TFR, 𝑒_0_, 𝑟, and 𝑇. All variables were Z-scored, and coefficients (𝛽) represent standardised effect sizes (**Table S6**). Model validity was assessed through residual analysis and collinearity checks (***Figure S 10*, Table S7**).

### Computational Environment

All analyses were performed in R (version 4.5.2). Package versions are recorded in *renv.lock* for reproducibility. Code and data are available at https://github.com/donertas-group/human_selection_shadow. The interactive selection shadow simulator is accessible at https://donertas-group.github.io/human_selection_shadow.

## Acknowledgements

HMD is funded by the Carl-Zeiss-Stiftung (P2021-00-007) and is a principal investigator in the Collaborative Research Centre CRC 1310 “Predictability in Evolution” funded by the Deutsche Forschungsgemeinschaft (DFG). The authors thank Hamit Izgi for helpful discussions on the manuscript.

## Supplementary Information

**Supplementary Figures**

**Figure S 1:**
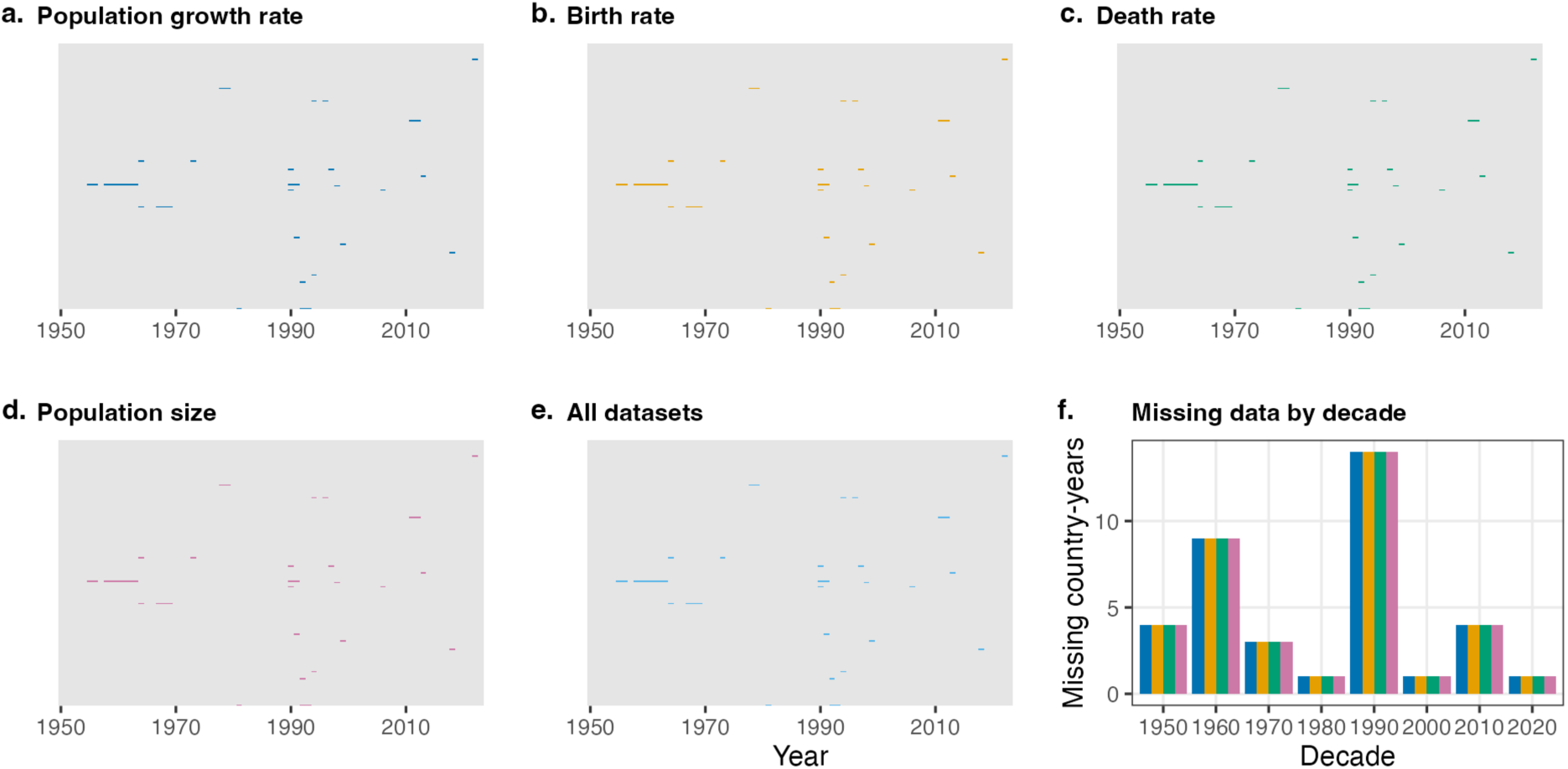
Data coverage across countries and years (1950–2023). **(a–e)** Heatmaps showing data availability for each variable: **(a)** population growth rate, **(b)** birth rate, **(c)** death rate, **(d)** population size, and **(e)** all datasets combined. Gray cells indicate data present; colored cells indicate excluded data. Exclusions reflect country-years with extreme population growth rates (|𝑟| > 10%) due to mass migration or wars (see Methods; **Table S2**). Each row represents a country/region; columns represent years. **(f)** Excluded country-years quantified by decade for each variable.

**Figure S 2:**
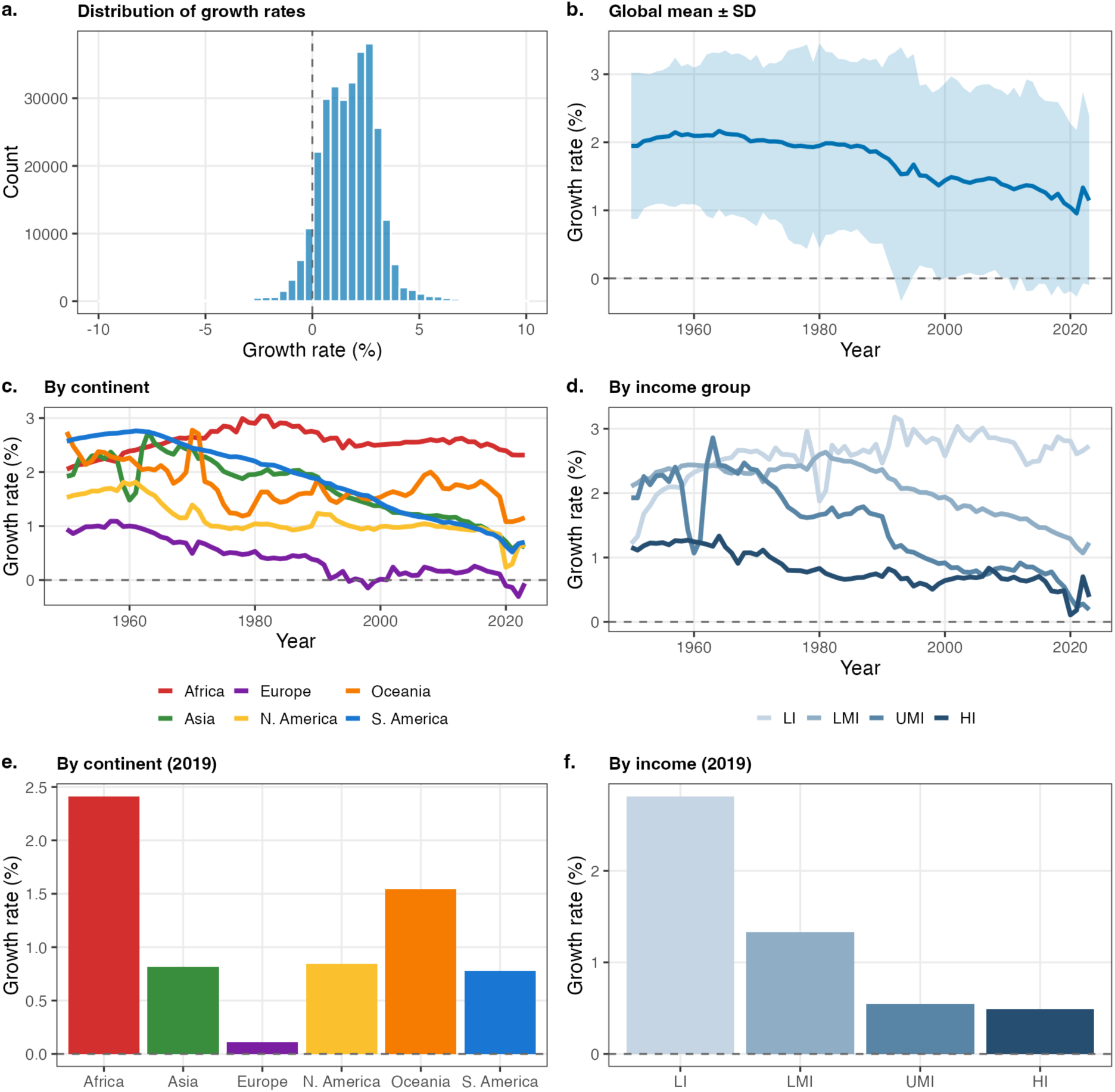
Population growth rate patterns across countries and time (1950–2023). **(a)** Distribution of growth rates (%) across all country-years. Dashed line marks zero growth. **(b)** Global mean ± SD over time; shaded region represents standard deviation. **(c)** Temporal trends by UN continent aggregate. **(d)** Temporal trends by World Bank income group aggregate. **(e)** Growth rate by continent in 2019. **(f)** Growth rate by income group in 2019.

**Figure S 3:**
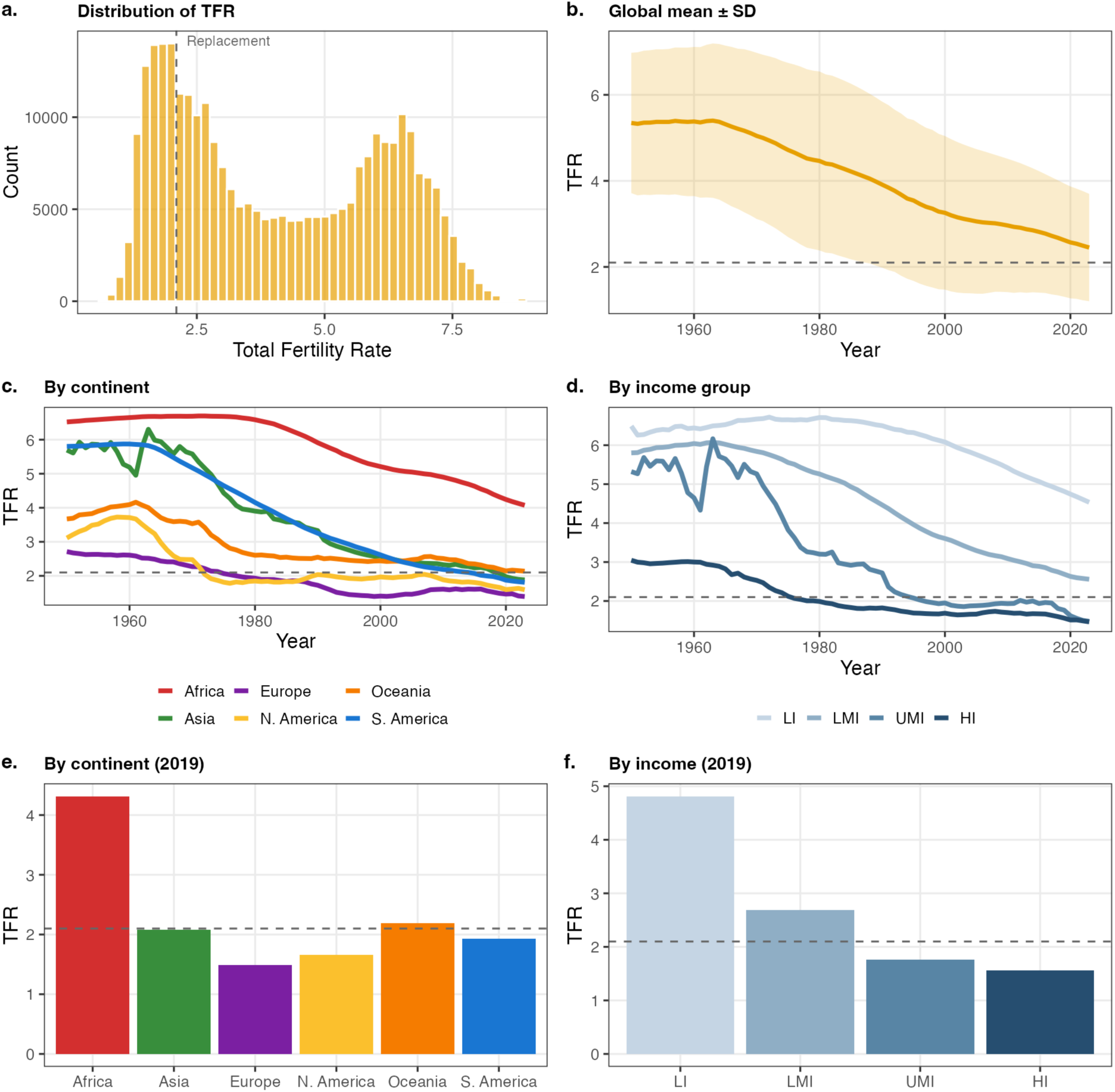
Total Fertility Rate (TFR) patterns across countries and time (1950–2023). **(a)** Distribution of TFR across all country-years. Dashed line marks replacement level (TFR = 2.1). **(b)** Global mean ± SD over time; shaded region represents standard deviation. **(c)** Temporal trends by UN continent aggregate; dashed line marks replacement level. **(d)** Temporal trends by World Bank income group aggregate. **(e)** TFR by continent in 2019. **(f)** TFR by income group in 2019.

**Figure S 4:**
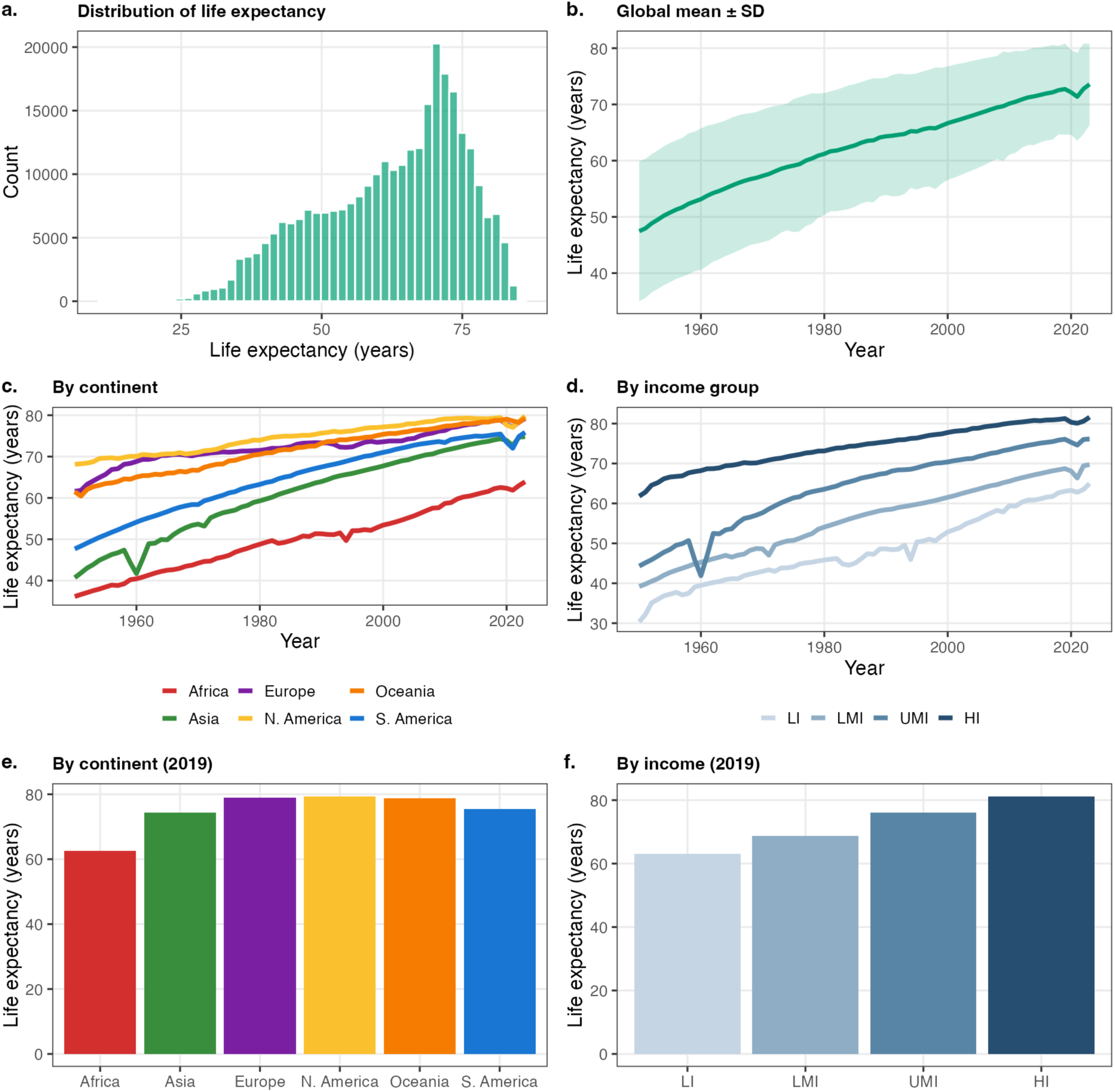
Life expectancy at birth (𝑒_0_) patterns across countries and time (1950–2023). **(a)** Distribution of 𝑒_0_ (years) across all country-years. **(b)** Global mean ± SD over time; shaded region represents standard deviation. **(c)** Temporal trends by UN continent aggregate. **(d)** Temporal trends by World Bank income group aggregate. **(e)** Life expectancy by continent in 2019. **(f)** Life expectancy by income group in 2019.

**Figure S 5:**
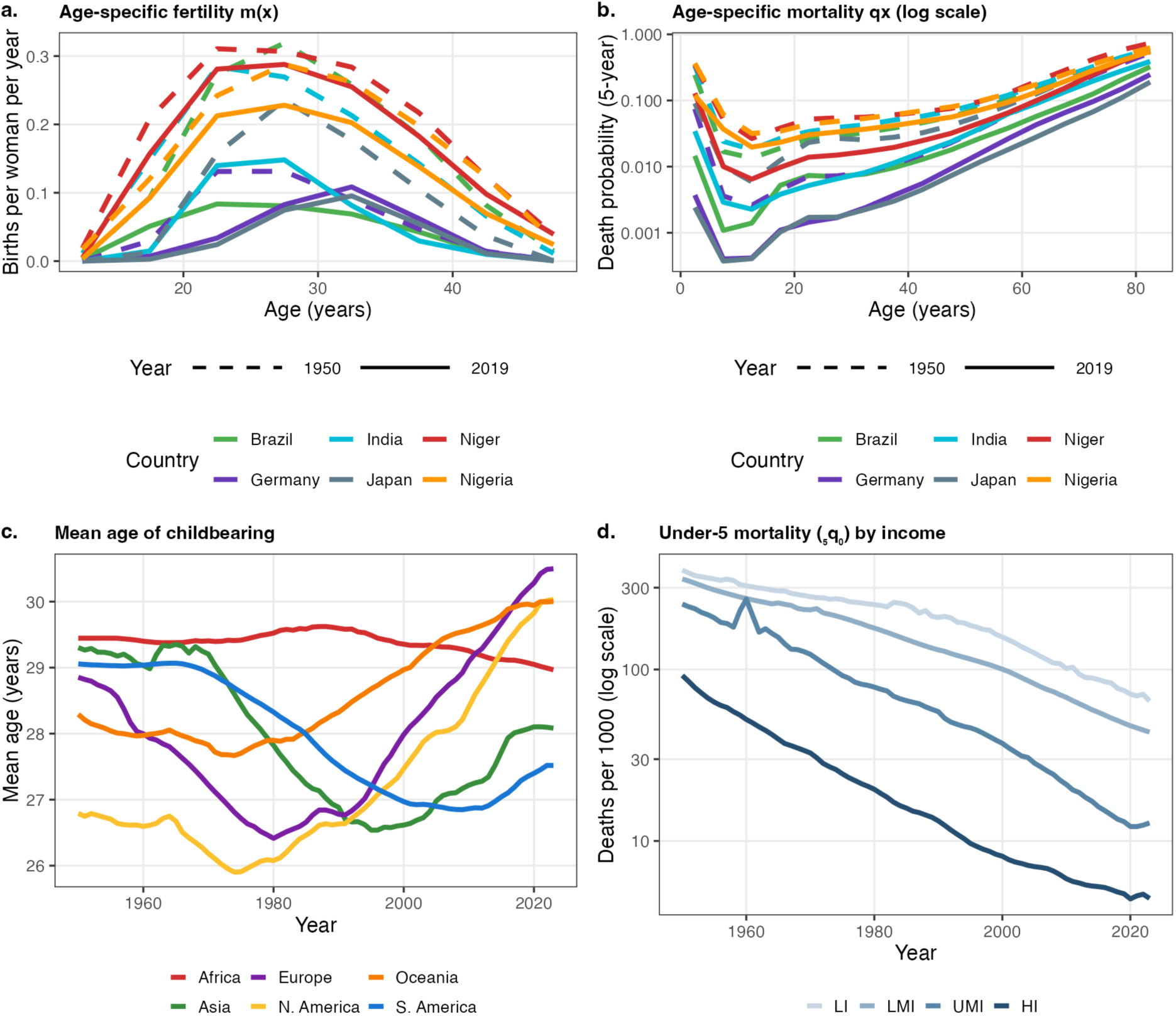
Age-specific demographic profiles. **(a)** Age-specific fertility rate 𝑚(𝑥) for six representative countries (Brazil, Germany, India, Japan, Niger, Nigeria) in 1950 (dashed) and 2019 (solid). **(b)** Age-specific mortality probability 𝑞_+_ (log scale) for the same countries and years. **(c)** Mean age of childbearing over time by UN continent aggregate. **(d)** Under-5 mortality (_5_𝑞_0_, deaths per 1000, log scale) by World Bank income group aggregate over time.

**Figure S 6:**
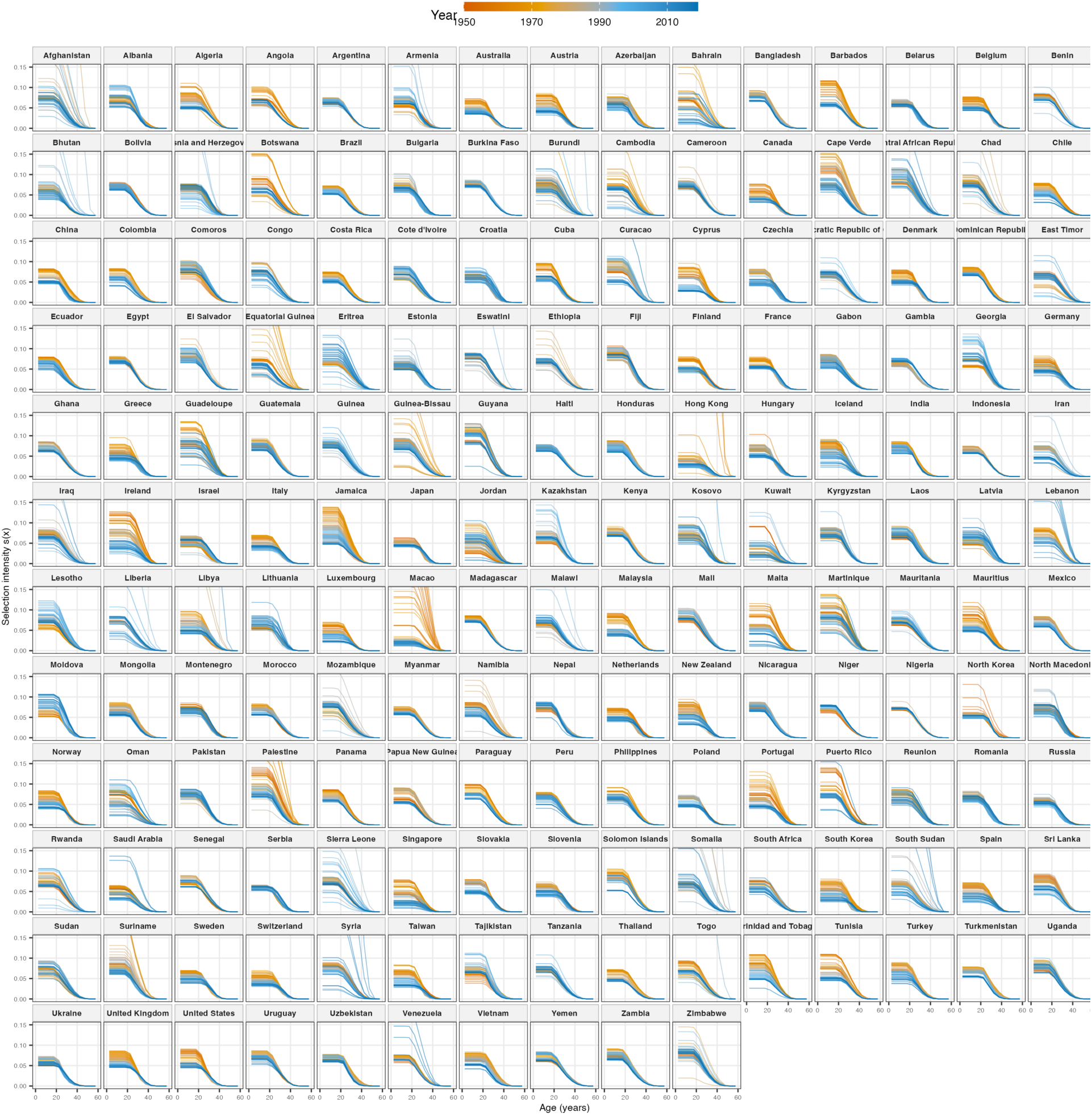
Global Atlas of Selection Shadows. Selection intensity curves 𝑠(𝑥) for all 165 nations and regions analyzed, colored by year from 1950 (orange) to 2019 (blue). Panels are ordered alphabetically. The grid reveals the global ubiquity of the “extension-dilution” effect: as populations transition (blue curves), the selection shadow consistently flattens, lowering 𝑠(0) while extending the age of selective relevance.

**Figure S 7:**
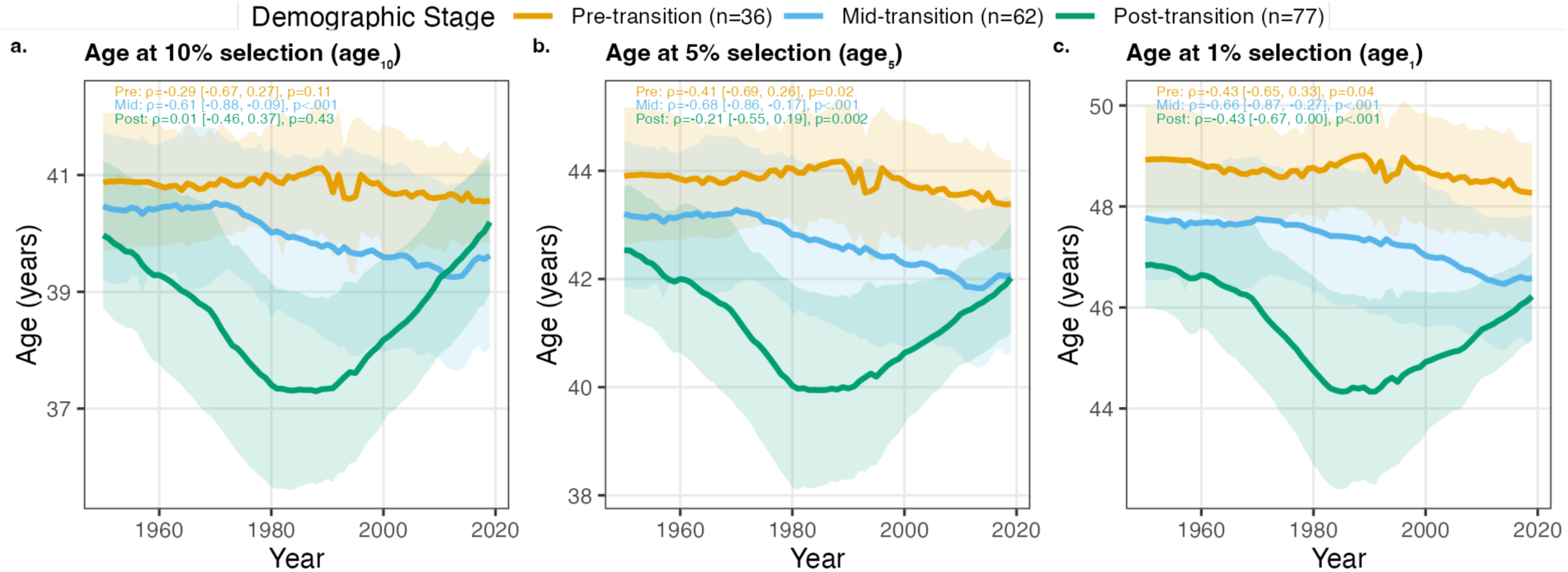
Temporal trends in selection thresholds.** Trends in the age at which selection intensity falls below **(a)** 10%, **(b)** 5%, and **(c)** 1% of baseline intensity. Post-transition populations (green) show a distinct “U-shaped” trajectory, where the selective window initially contracted (1950–1980) before expanding again in recent decades, likely driven by the stabilization of fertility timing and continued mortality compression.

**Figure S 8:**
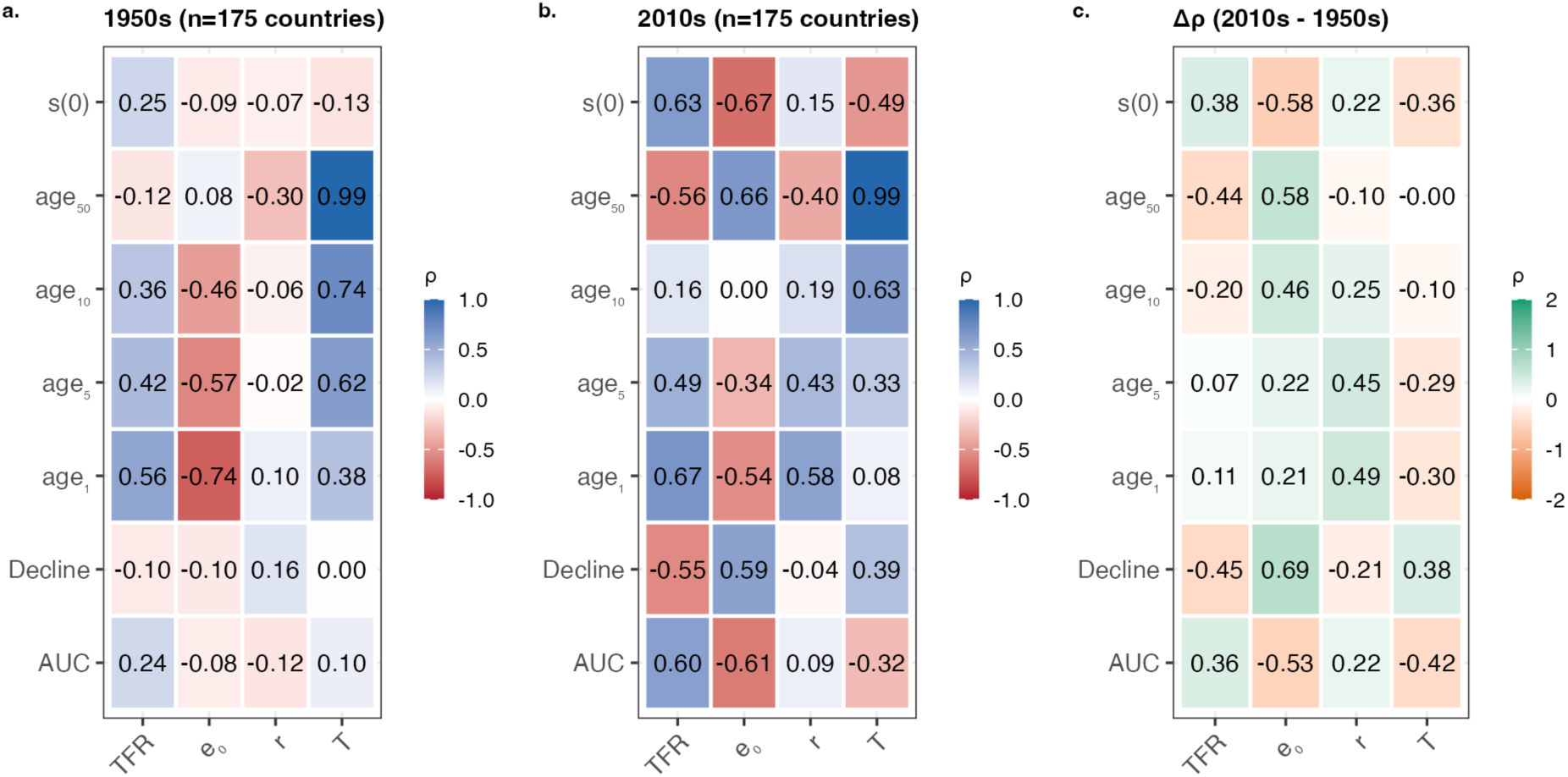
Restructuring of demographic correlations.** Correlation matrices (Spearman’s 𝜌) between selection metrics (𝑠(0), age_50%_, etc.) and demographic drivers (TFR, 𝑒_0_, 𝑟, 𝑇) for **(a)** the 1950s and **(b)** the 2010s. **(c)** The difference 𝛥𝜌 (2010s - 1950s). Green indicates the correlation became more positive; orange indicates it became more negative.

**Figure S 9:**
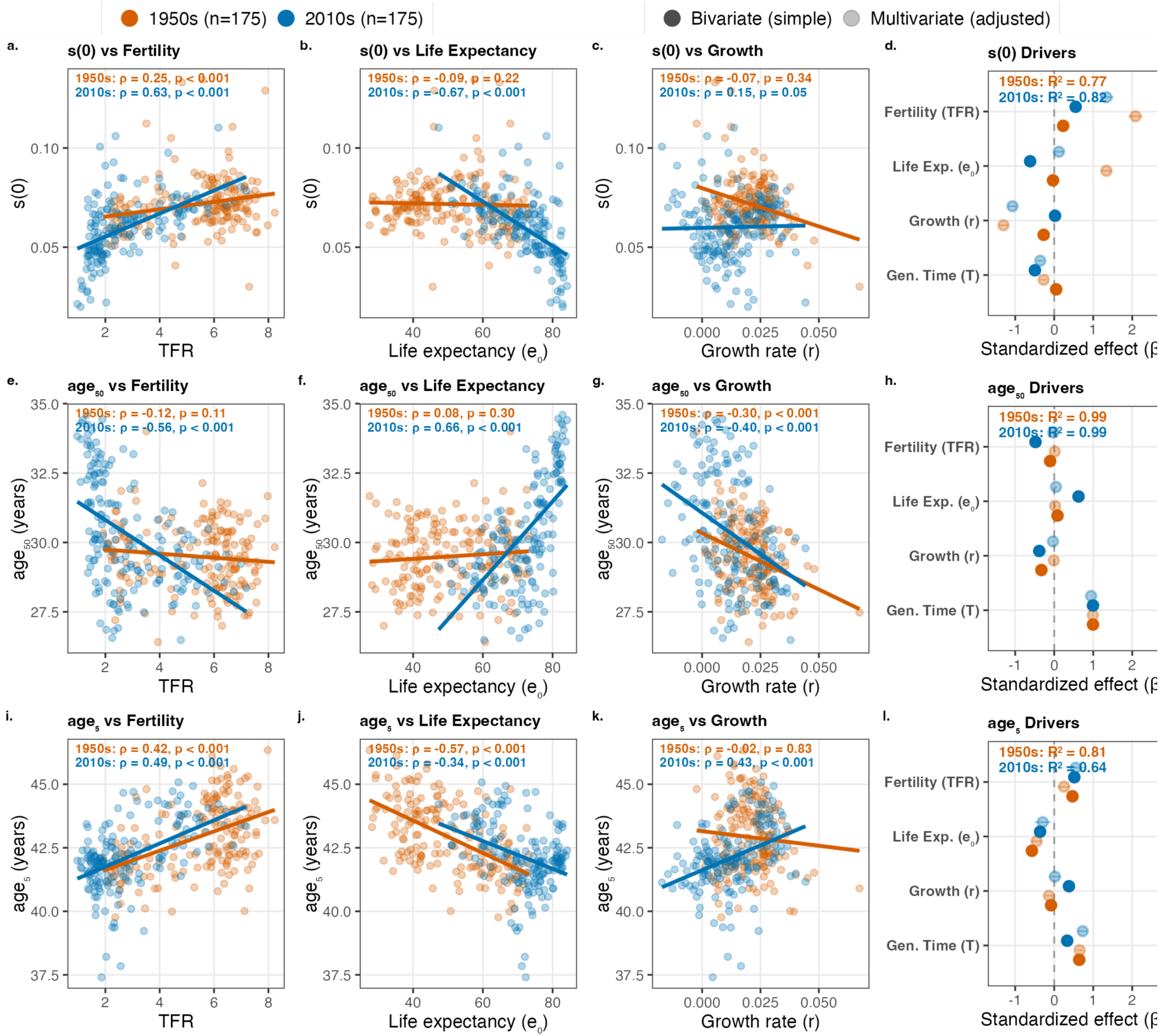
Demographic determinants of selection metrics. Scatter plots and standardized regression coefficients showing relationships between selection metrics and demographic drivers, comparing 1950s (orange) and 2010s (blue). **(a–d)** 𝑠(0) relationships: **(a)** vs. TFR, **(b)** vs. life expectancy, **(c)** vs. growth rate, **(d)** coefficient plot. **(e–h)** age_50_ relationships. **(i–l)** age_5_ relationships. Each scatter plot shows Spearman correlations with explicit p-values. Coefficient plots display both bivariate (solid) and multivariate (faded) standardized effects with 95% CIs.

**Figure S 10:**
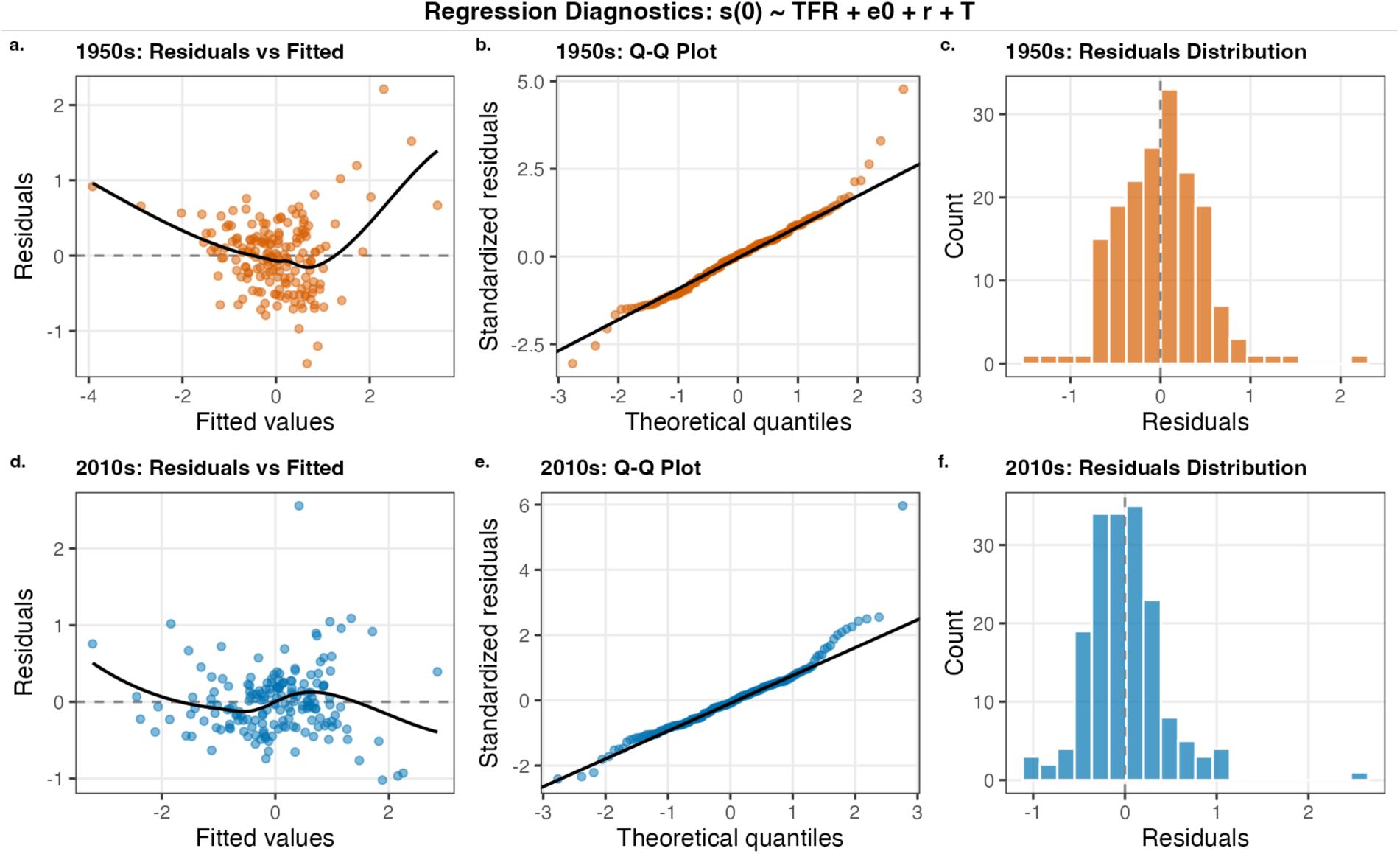
Regression Diagnostics. Diagnostic plots for the multivariate regression models predicting 𝑠(0) (𝑠(0) ∼ 𝑇𝐹𝑅 + 𝑒_0_ + 𝑟 + 𝑇). **(a-c)** 1950s diagnostics: Residuals vs. Fitted, Q-Q plot, and Residual Histogram. **(d-f)** 2010s diagnostics.

## Supplementary Tables

**Table S1. Excluded small populations.** List of countries and territories excluded from the analysis due to small population size (total population < 100,000). For each entity, the table reports the number of excluded country-years and the year range affected. Small populations were excluded to minimise noise arising from demographic stochasticity and small-number artefacts in age-specific rate calculations.

**Table S2. Excluded extreme growth events.** List of country-year observations excluded due to extreme population growth volatility (|r| > 10%). These observations represent instances of massive structural shocks, where instantaneous growth rates do not reflect biological fitness and violate the stable population assumption underlying the selection shadow calculation.

**Table S3. Excluded artifact observations.** List of country-year observations excluded from temporal trend analyses and regression modelling due to biologically implausible selection values (𝑠(0) ≤ 0 or 𝑠(0) ≥ 0.20). For each observation, the table reports: country, year, demographic stage, 𝑠(0), age_50%_, 𝑅_0_, and generation time (𝑇).

**Table S4. Country-year selection metrics.** Complete dataset of selection metrics for all 175 countries and 16 aggregate regions from 1950 to 2019. For each country-year observation, the table includes: selection intensity at birth (𝑠(0)), age at 50% selection (age_50%_), selection intensity at the half-life point, ages at 10%, 5%, and 1% selection thresholds, selection intensity at fixed ages (20, 40, 50, 60), area under the curve (AUC), decline rate, generation time (𝑇), net reproductive rate (𝑅_0_), and demographic stage classification.

**Table S5. Temporal summaries by decade.** Decadal summary statistics for selection metrics across all populations. For each decade (1950s–2010s), the table reports: number of country-years and countries, mean, standard deviation, median, minimum, and maximum values for 𝑠(0), age_50%_, AUC, decline rate, generation time (𝑇), and net reproductive rate (𝑅_0_).

**Table S6. Regression coefficients.** Standardised regression coefficients from multivariate models predicting selection metrics (𝑠(0), age_50%_, age_5%_) from demographic drivers (TFR, 𝑒_0_, 𝑟, 𝑇). For each period (1950s, 2010s) and response variable, the table reports: standardised coefficient, standard error, p-value, significance level, and model 𝑅^+^.

**Table S7. Regression diagnostics.** Model fit statistics for multivariate regression models. For each period and response variable, the table reports: sample size (𝑛), 𝑅^+^, adjusted 𝑅^+^, root mean squared error (RMSE), mean absolute error (MAE), AIC, BIC, Shapiro-Wilk test statistic and p-value for residual normality, and residual range.

## Notes

### Competing Interest Statement

The authors have declared no competing interest.

